# Bidirectional selection for body weight on standing genetic variation in a chicken model

**DOI:** 10.1101/469049

**Authors:** Mette Lillie, Christa F. Honaker, Paul B. Siegel, Örjan Carlborg

**Author notes:** Current affiliations: Department of Biological and Environmental Sciences, University of Gothenburg, Gothenburg, Sweden and Department of Ecology and Genetics, Uppsala University, Uppsala, Sweden (Mette Lillie). **Corresponding author information:**, **Name:** Mette Lillie, **Email address:**, **Mailing address:** University of Gothenburg, Dept for Biological and Environmental Sciences, Mette Lillie, PO Box 463, SE 405 30 Göteborg. Data Availability Statement: Pooled genome data generated for this study will be deposited in the Short Read Archive and accession numbers will be included in the manuscript. Supplementary Material (Figures S1, S2, and S3 and Tables S1, and S2) has been deposited on fig**share**.

## Abstract

Experimental populations of model organisms provide valuable opportunities to unravel the genomic impact of selection in a controlled system. The Virginia body weight chicken lines represent a unique resource to investigate signatures of selection in a system where long-term, single-trait, bidirectional selection has been carried out for more than 60 generations. Using pooled genome resequencing of paired generations of these lines, we reveal the within and between-line genomic signatures of selection. At 55 generations of divergent selection, 14.2% of the genome showed extreme differentiation between the selected lines were contained within 395 genomic regions. The lines often displayed a duality of the sweep signatures: an extended region of homozygosity in one line, in contrast to mosaic pattern of heterozygosity in the other line. These haplotype mosaics consist of short, distinct haploblocks of variable between-line divergence. Formed during what probably was a complex history of bottlenecks, inbreeding, and introgressions, these mosaics represent the standing genetic variation available at the onset of selection in the founder population. Selection on standing genetic variation can thus result in different signatures depending on the intensity and direction of selection.

## Introduction

Initial thinking on how adaptive processes shape the genome was modeled by Maynard Smith and Haigh (1974), who demonstrated that a beneficial mutation favored by natural selection will increase in frequency within a population. Linkage disequilibrium in the flanking region of a selected allele will result in a characteristic valley of diversity around the selected variant, known as a hard selective sweep (Kaplan et al. 1989; Stephan et al. 1992). Selection on recessive variants, variants contributing to the standing genetic variation in a population, and partial sweeps (Hermisson and Pennings 2005; Teshima and Przeworski 2006) typically have weaker effects on linked sites, but empirical studies suggest that such selection is the most abundant mode of adaptation in recent evolution in both *Drosophila melanogaster* (Garud et al. 2015) and humans (Pritchard et al. 2010; Hernandez et al. 2011; Schrider and Kern 2017). Furthermore, polygenic adaptation describes how selection acts on standing genetic variation across the many loci contributing to a quantitative polygenic trait leading to a new phenotypic optimum by way of modest allele frequency changes across these loci (Pritchard and Di Rienzo 2010; Pritchard et al. 2010). Although this mode of adaptation would respond very rapidly to changes in the selective environment, it would not necessarily lead to fixation for any one variant (Pritchard et al. 2010).

Modes of adaptation are not mutually exclusive, and the genomic signature that results will be dependent upon factors such as the genetic architecture of the trait, standing genetic variation available within this architecture, effective sizes of standing haplotypes, and population demography. By exploiting population genetic signals, researchers are increasingly able to detect the underlying modes of selection, from initial sweep scans that identify valleys of low diversity resulting from hard sweeps, to various recent developments to detect and differentiate between soft and hard sweeps (Berg and Coop 2014; Garud et al. 2015; Schrider and Kern 2017). Whereas previous studies have attempted to uncover selection throughout human history (Coop et al. 2009; Schrider and Kern 2017), much can be learned from research with model organisms, such as selection in experimental populations of *Drosophila* (Burke et al. 2010) or mice (Chan et al. 2012). In particular, long-term selection experiments have well-defined population histories, likely have stronger selection signatures in the genome due to an isolation of the trait under selection, and allow breeding of crosses to test for adaptive trait associations to candidate sweeps. The Virginia body weight chicken lines, whose history of long-term, single-trait, bi-directional selection from a common founder population affords us the opportunity to dissect the genomic selective-sweep signatures of strongly selected loci (Figure 1).

**Figure 1.**
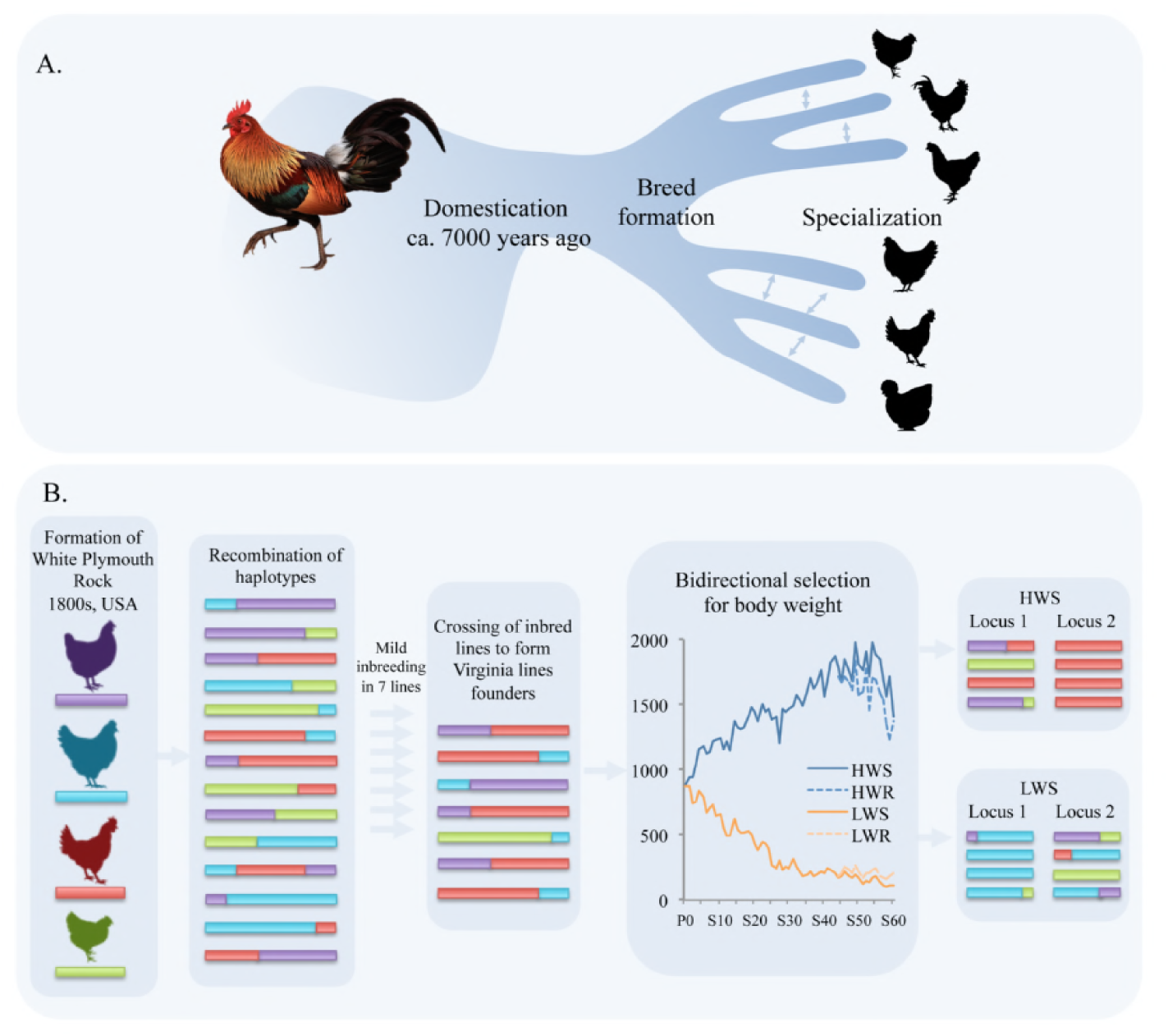
The evolutionary history of the Virginia body weight chicken lines makes them a powerful model to dissect the genomic signature of selective-sweeps contributing to polygenic adaptation. A. After domestication approximately 7,000 years ago, chicken breeds were gradually established with specialization for production traits, such as egg-laying, meat, or dual-purpose. Breeds with novel characteristics such as duration of crows, length of tail feathers, or feather patterns were also established.
B. The White Plymouth Rock was formed in America in the 1800s from crossing of several established breeds (Dohner 2001). During this breed development, divergent haplotypes contributed from established breeds would recombine along the genome. Founders for the Virginia lines were generated via crossing of seven partially inbred White Plymouth Rock lines, which would have constrained the standing genomic diversity. From 1957, bidirectional selection for 8-week body weight was carried out to form the high (HWS) and low (LWS) Virginia body weight lines. The HWS and LWS are highly differentiated for a large portion of the genome, including those with known associations to the selected trait.

Here, we utilize the Virginia body weight chicken lines to investigate the genomic impact of selection. These lines were established in 1957 from partially inbred White Plymouth Rock chickens (WPR). Two lines (high weight selected: HWS; low weight selected: LWS) were selectively bred for divergent body weight at 8 weeks of age with the breeding regime structured to minimize inbreeding and minimize the stochastic fixation of alleles that could affect small breeding populations (Siegel 1962; Marquez et al. 2010). After 55 generations of divergent selection, a 15-fold difference in body weight exists between the lines (Jambui et al. 2017). With its well-defined population history, bi-directional single trait selection regime, and well-defined polygenic architecture of the adaptive trait, the Virginia body weight chicken lines represent an invaluable resource to investigate the genetic and genomic signatures resulting from the polygenic adaptation that has driven the long-term selection response.

Previous research has demonstrated that standing genetic variations in many loci from the founder population of the Virginia lines contribute to the observable difference in body weight. Sweep scans using individual genotypes from a 60k SNP-chip have revealed numerous regions of differentiation between the selected lines (Johansson et al. 2010; Pettersson et al. 2013), of which some were associated with body weight and contributed to selection response using available standing genetic variation (Sheng et al. 2015). Recently, 20 independent associations to 8-week body weight were confirmed using genotypic and phenotypic data from an F_15_ intercross (Zan et al. 2017) and pooled genome re-sequencing data from the HWS and LWS lines (Lillie et al. 2017a). The accumulation of evidence suggests that high allelic complexities across associated loci within the Virginia body weight lines involve multiple alleles, linked loci, and epistasis contributing to the polygenic architecture of body weight (Lillie et al. 2017a; Zan et al. 2017).

We used pooled genome resequencing to characterize the genomic signatures of these highly differentiated regions between the bidirectionally-selected lines: the putative selective sweeps. Between-line differentiation was seldom due to complete fixation of different haplotypes in the lines. Instead, most selective sweeps resulted from extended runs of homozygosity in one line, contrasting to persistence of heterozygosity in the other. Typically, this heterozygous region was comprised of multiple, distinct regions, with variable diversity, and between-line divergence. The directions of these relationships do not suggest any line bias.

These empirical observations and our knowledge about the history of these populations imply that the duality of these signatures reflect positive selection for one large-effect haplotype in one line while in the other line negative selection would remove this haplotype, allowing other haplotypes to continue to segregate. Mosaic haplotype structures within these segregating regions reflect standing variation in the founder population, likely resulting from ancestral haplotype recombination along a history of bottlenecking, inbreeding, and crossbreeding.

## Materials and Methods

### Virginia body weight chicken lines

All animal procedures were carried out by experienced handlers and in accordance with the Virginia Tech Animal Care Committee animal use protocols (IACUC-15-136). The Virginia body weight lines were formed from a founder population resulting from crossing seven lines originating in 1949 that had undergone mild inbreeding. Established in 1957, bidirectional selection for body weight at 8 weeks of age was initiated to produce the closed selected lines: high weight selected (HWS) and low weight selected (LWS) (Siegel 1962). Breeding focused on a response to selection, while attempting to minimize inbreeding (Marquez et al. 2010).

All generations were hatched in the same incubators and reared in the same pens on the same diet. Relaxed sublines for both HWS and LWS were produced from selected generation 44, and are referred to as high weight relaxed (HWR) and low weight relaxed (LWR) (Dunnington et al. 2013). Pooled semen was used to produce each generation of relaxed lines.

### Sequencing and genome alignments

DNA for the genomic analyses was prepared from blood samples collected from 9-30 individuals from each line and pooled in equimolar ratios prior to library construction. Genome sequencing library construction and sequencing was carried out by SciLifeLab (Uppsala, Sweden) using two lanes on an Illumina Hiseq 2500. Pooled genome data generated for this study are available via SRA (bioproject: TBA; SRA accession numbers: TBA). Reads were aligned to the *Gallus gallus* genome (Galgal5; INSDC Assembly GCA_000002315.3, Dec 2015) using BWA (Li and Durbin 2009). Genomes were sorted and duplicates were marked and removed with Picard (v1.92; http://picard.sourceforge.net). GATK (v3.3.0; McKenna et al. 2010) was used for realignment around indels. GATK UnifiedGenotyper was used to generate allele calls at all sites (option: emit all sites) and with ploidy = 30 (18 for LWS generation 50 as only 9 individuals went into this pool) to account for the pooled genome sample. Sites were filtered to only include those with >10 and <100 reads, wherefrom allele frequency, heterozygosity, and pairwise *F_ST_* between all populations were calculated. Samtools (Li et al. 2009; v1.1; Li 2011) was used to generate mpileup files for PoPoolation2 (v1.201; Kofler et al. 2011), which was used to calculate *F_ST_* over 1000 bp sliding windows with 50% overlaps between the population samples using the Karlsson et al. (2007) method, with minimum count 3, minimum coverage 10, maximum coverage 100, and minimum coverage fraction 1. Genome alignments were visualized in IGV (v2.3.52; Robinson et al. 2011; Thorvaldsdottir et al. 2013).

### Differentiated regions

*F_ST_* cutoff = 0.953 (95% percentile of *F_ST_* values in generation 55). Windows with *F_ST_* greater than this cutoff were used to limit the number of candidate regions due to drift. Windows with *F_ST_* values above this cutoff were clustered into differentiated regions when they were less than 100 kb from one another (custom R scripts). Clusters with less than 2 SNPs or less than 100 kb were removed from the dataset to retain only the stronger candidate regions. Mean and median heterozygosity were calculated for each line within each differentiated region. We used the Variant Effect Predictor (VEP) (McLaren et al. 2016) available from Ensembl (Aken et al. 2017) to investigate potential functionality of candidate alleles. Genomic / haplotypic signatures within regions of interest were visualized by adjusting allele frequencies as used in (Lillie et al. 2017b) (similar to the allele polarization step in the haplotype-block reconstruction approach used by Franssen et al. 2017) to the generation of lowest complexity, then plotted using custom R scripts. Pooled genome data generated for this study are available via SRA (bioproject: TBA; SRA accession numbers: TBA).

## Results

From clusters of high differentiation (*F_ST_* > 0.953; Figure 2), we identified 395 differentiated regions between the HWS and LWS lines in selected generation 55 (Lillie et al. 2017a). These regions covered a total of 174.5 Mb, or 14.2% of the genome, which is an increase from the 244 differentiated regions identified in selected generation 40 (99.6 Mb or 8.1% of the genome; Figure S1)(Lillie et al. 2017a).

**Figure 2.**
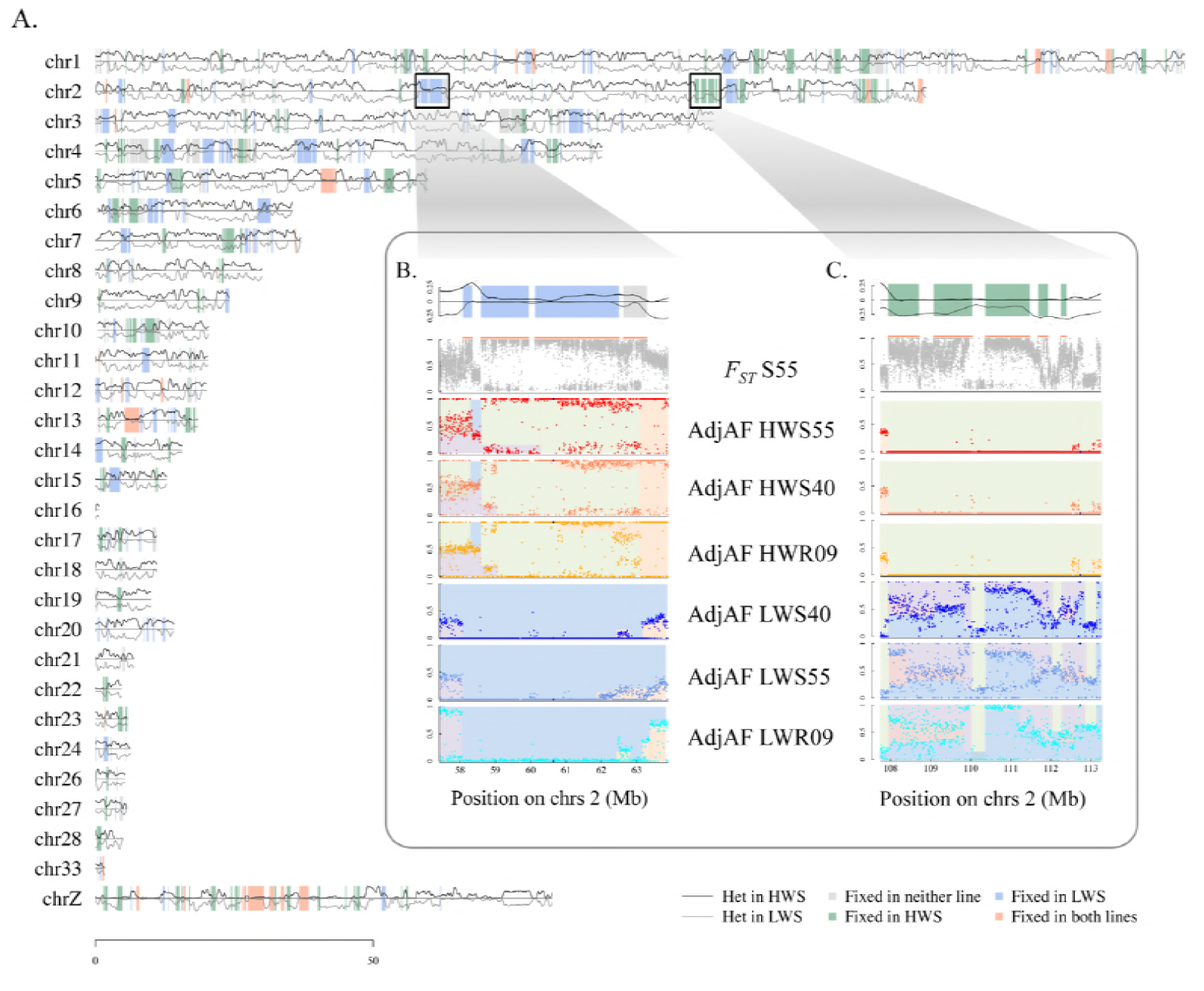
Heterozygosity across chromosomes of divergently selected Virginia body weight chicken lines. A. Heterozygosity in generation 55 for HWS (black line above axis) and LWS (grey line above axis) presented across the chicken chromosomes within the differentiated regions shaded using color code: grey where neither line is fixed; green where only HWS is fixed; blue where only LWS is fixed within the differentiated region; red where both lines are fixed.
B. *F_ST_* and heterozygosity patterns across chromosome 2:58-63 Mb for selected generations 55 and 40 and relaxed generation 9.
C. *F_ST_* and heterozygosity patterns across chromosome 2:108-113 Mb for selected generations 55 and 40 and relaxed generation 9. Panels within B and C insets: Panel 1: Detail of heterozygosity trace from chromosome map. Panel 2: Mean *F_ST_* within 1 kb windows indicated with grey points; region of differentiation indicated with orange line above *F_ST_* plot. Panel 3-8: Mean adjusted allele frequency of 5kb windows in HWS_55_/HWS_40_/HWR_09_/LWS_55_/LWS_40_/ LWR_09_. Colors within the plots highlight the runs of adjusted allele frequencies that contribute to different haploblocks.

### Candidate selective sweeps are often polymorphic in one of the selected lines

Few regions of differentiation showed fixation in both lines; rather, it was often the case that there was fixation across an extended region in one line, while many nucleotide positions in the region still segregated in the other (Figure 3; S2 Figure). This demonstrated that, while one extended haplotype was fixed in one line, multiple haplotypes continue to segregate in the other. Fixation was as common in HWS as LWS, a trend that extended to regions with confirmed associations for the selected trait, 8-week body weight (Figure 3; S2 Figure) (Zan et al. 2017).

**Figure 3.**
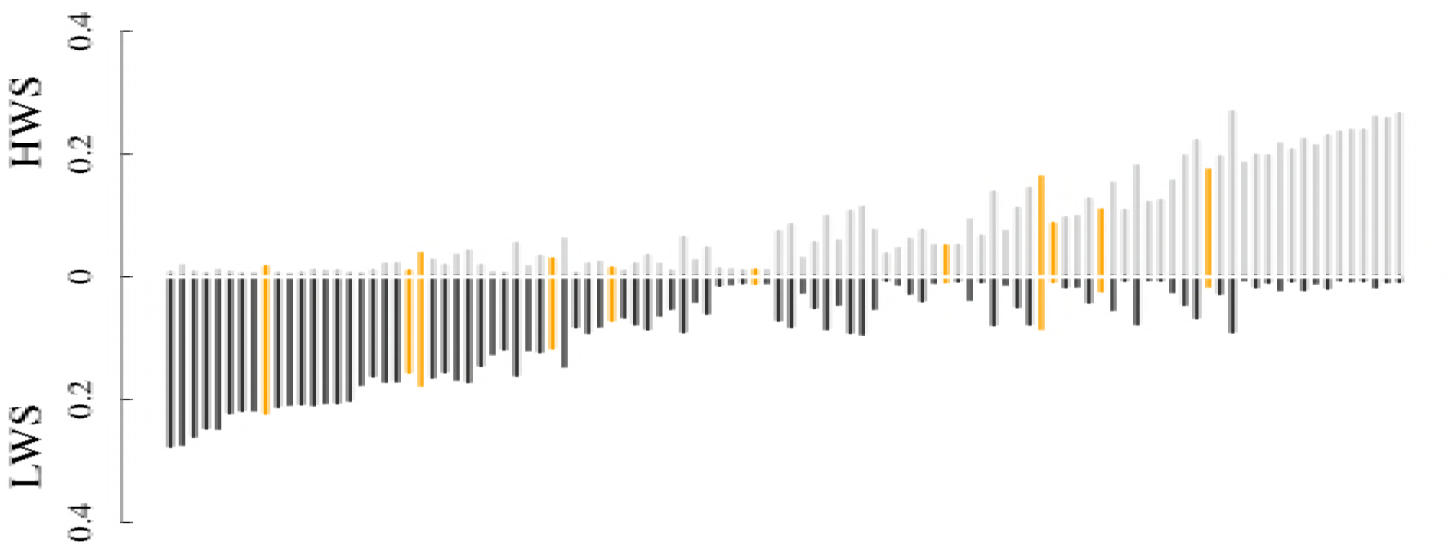
Candidate selective sweeps identified in the Virginia body weight chicken lines are often polymorphic in one, but not the other, selected line. Comparisons of mean heterozygosity remaining in HWS (above axis; grey) and LWS (below axis; black) at generation 55 within differentiated regions greater than 0.5 Mb in length. Differentiated regions overlapping known associations with 8-week body weight (Zan et al. 2017) are indicated in orange.

### Mosaic haplotypes in the growth1 QTL

To illustrate the genomic mosaicism observed in many differentiated regions, we investigated the *growth1* QTL in detail (Jacobsson et al. 2005; Zan et al. 2017). A long region of divergence was observed between 169.3 Mb and 173.7 Mb (in total 4.4 Mb), where a single extended haplotype was close to fixation in LWS by generation 40, whereas the pattern of polymorphism in the HWS suggest that multiple haplotypes segregate in this line (Figure 4). Approximately 28% of the nucleotide positions within this region were highly divergent between HWS and LWS at generation 55. In total, 10,148 from 36,934 sites had differences between the lines in allele frequency that were greater than 0.9. This level of divergence implies that the long haplotype fixed in LWS is unlikely to be present in the HWS at generation 55, likely because it has been selected out.

**Figure 4.**
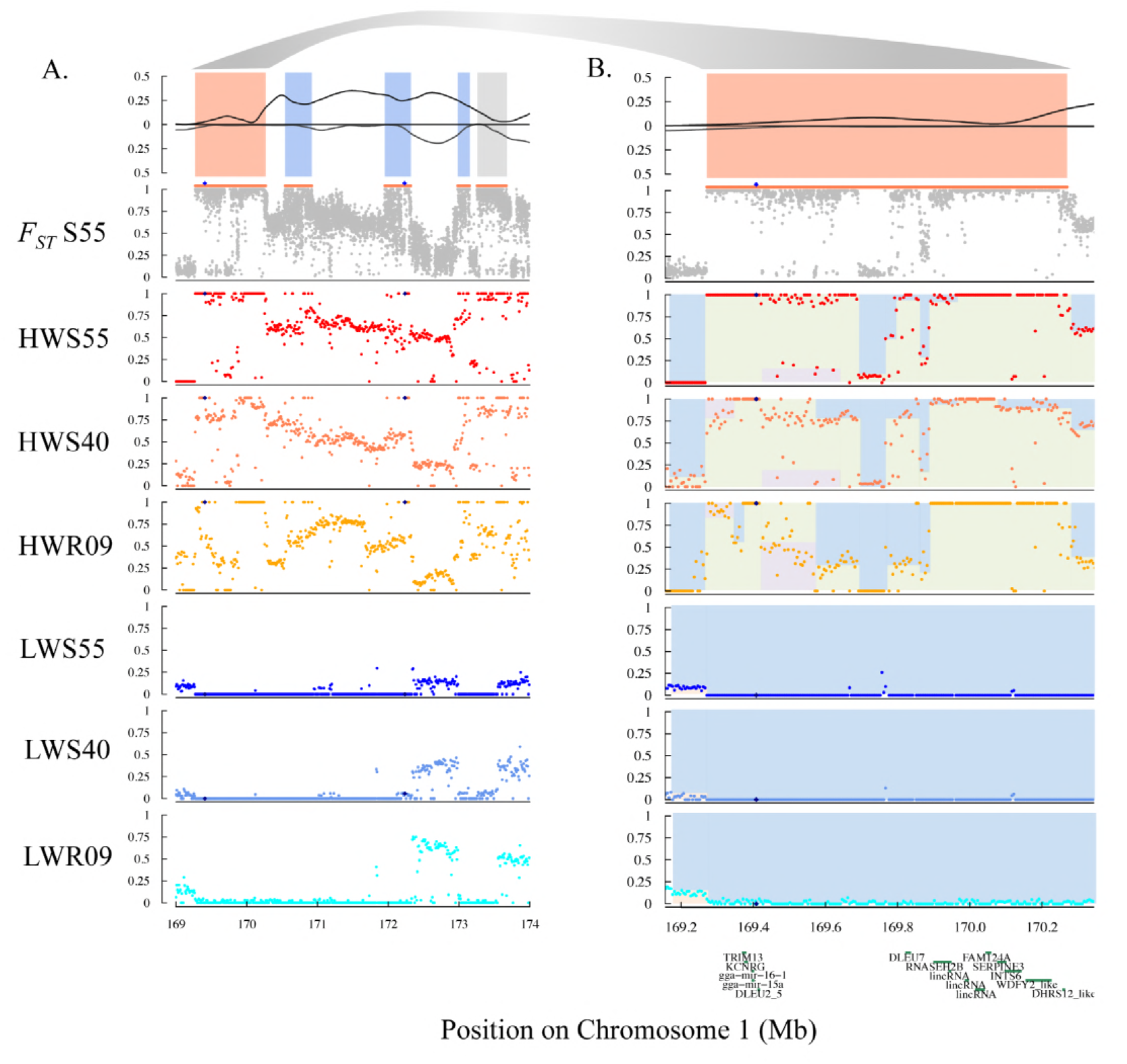
A. *F_ST_* and heterozygosity patterns across chromosome 1:169-174 Mb for selected generations 55 and 40 and relaxed generation 9 of the Virginia body weight chicken lines. B. *F_ST_* and heterozygosity patterns across chromosome 1:169.2-170.3 Mb for selected generations 55 and 40 and relaxed generation 9 with gene positions. Panel 1: Heterozygosity. Panel 2: Mean *F_ST_* within 1 kb windows (grey points) in selected generation 55; defined region of differentiation indicated with orange line above *F_ST_* plot. Panel 3-8: Mean adjusted allele frequency of 5kb windows in HWS_55_ (panel 3), HWS_40_ (panel 4), HWR_09_ (panel 5), LWS_55_ (panel 6), LWS_40_ (panel 7), LWR_09_ (panel 8). Colors within the plots are used to highlight the runs of adjusted allele frequencies that contribute to different haploblocks.

We checked for the presence of a known insertion that has been previously associated with increased body weight in other chicken populations (Jia et al. 2016). By examining the soft-clipped reads (S3 Figure), we observed that this variant also segregated in our population within the short region fixed for divergent haplotypes in both LWS and HWS (approximately 50 kb in length; GGA1: 169,370,000-169,420,000). Contrary to the expectation that this insertion would dominate in the HWS line, it was instead present on the long haplotype fixed by generation 40 in LWS, and entirely absent in HWS. Furthermore, this insertion is linked to the LWS allele of SNP marker rs14916997, which was associated with low body weight (Zan et al. 2017).

To identify candidate linkages and thus other potential functional variants, the *growth1* QTL was evaluated for further sub-haplotype fixations. In the nearby region (GGA1: 169,950,000-170,220,000), the HWS and LWS were also fixed by generation 55. Bounded by these two highly differentiated regions are 15 annotated genes (S1 Table), with 8 missense mutations (S2 Table). One missense mutation with predicted (SIFT) deleterious effect was present in HWS, located within the sixth exon of Ribonuclease H2 Subunit B gene (RNASEH2B), as well as the small (18 bp) deletion in LWS at GGA1: 169,407,811, which was absent from HWS.

## Discussion

### Candidate selective sweeps were often polymorphic in one of the selected lines

Within highly differentiated candidate selective sweep regions, we observed that an extended haplotype was often fixed in one line while multiple haplotypes continued to segregate in the other, thus resembling a duality in these sweep signatures. These patterns are likely to be shaped by the intensity of selection on the locus, as well as the haplotype frequencies that were present in the base population. For example, haplotypes with equal but opposite effects present at the same frequencies at the onset of selection would be expected to result in relatively equal lengths of fixation (homozygosity) in both lines, and thus a high *F_ST_*. This pattern was observed only a few times (Figure 3), and thus appears to have been a relatively rare event. More likely is that the haplotypes in the founder population had different effect sizes and were present at different frequencies when selection was initiated to develop the high and low body weight chicken lines. The long runs on homozygosity found only in one line within regions of differentiation therefore likely represent the haplotypes with the largest effect, with the signature possibly being amplified by being at a low frequency at the onset of selection. These patterns suggest that the effect size of functional variants, as well as founding haplotype frequencies, are likely to have influenced the selective signature in the genome of the Virginia body weight chicken lines.

### Mosaic haplotypes

Commencing with the domestication of the junglefowl, the history of the Virginia body weight chicken lines is characterized by multiple factors including population bottlenecks, inbreeding, outbreeding, and changes in selection pressures (Figure 1). Domestication of the chicken began roughly 7,000 years ago, predominately from red junglefowl, *Gallus gallus* (West and Zhou 1988; Sawai et al. 2010), but potentially also with contributions from *Gallus sonneratii* (grey) and *Gallus layafetii* (Sri Lankan) (Eriksson et al. 2008; Groeneveld et al. 2010; Tixier-Boichard et al. 2011). With subsequent distribution of chickens via human dispersal and trade, local breed formation would subject established chicken populations to genetic drift and selection to suit the environment and human-imposed selection for breed standards and production traits (Lyimo et al. 2014). Regardless of expectations that founder effects, bottlenecks, and selection would impact genetic diversity, nucleotide diversity and substitution rates were comparable for red jungle fowl and domestic chickens. This implies that domestication did not result overall in a substantial genome-wide loss of diversity in the species and had only a minor effect in an evolutionary context (Tixier-Boichard et al. 2011). The Plymouth Rock is a relatively modern breed; first exhibited in mid-19^th^ century as the pioneer barred variety from which the White Plymouth Rock variety was later developed (Smith 1921). The exact contributions of breeds into Plymouth Rock has been debated, but likely originated from crosses of Dominique, Black Java, and Cochin (Smith 1921), with potential input from Brahma (Howard 1907). Thus, recombination of divergent haplotypes from European (Dominique) and Asian (Java, Cochin, and Brahma) populations occurred while producing the WPR breed, forming the genetic substrate available for artificial selection (Figure 1). During the several decades of WPR breeding that preceded the initiation of the Virginia body weight chicken lines, further recombinant haplotypes are likely to have arisen, ultimately producing finely grained mosaics of the chromosomes entering the founders of the Virginia body weight lines. It is on this standing variation which the bi-directional selection experiment has acted. Short haplotype mosaics within the longer selective sweep regions were revealed when comparing the HWS and LWS genomes. The *growth1* QTL region on chromosome 1 is an example, illustrating the influence of the population history on the fine-grained selective sweep signatures observed across the genome.

### Mosaic haplotypes in the growth1 QTL

The *growth1* QTL was first defined in a microsatellite-based analysis of an F_2_ cross between the HWS and LWS, with a length of almost 110 cM (Jacobsson et al. 2005). This QTL was confirmed in SNP-based analyses of the F_2_ (Wahlberg et al. 2009) and fine mapped in the F_2_-F_8_ (Besnier et al. 2011; Brandt et al. 2016) and the F_15_ generation of the Advanced Intercross Line (Zan et al. 2017). In this latest study, two associations were found within *growth1*, with the SNP rs14916997 (GGA1: 169,408,309) having the strongest association (Zan et al. 2017). From the pooled genome data, it is evident that a major haplotype, between 169.3 Mb and 173.7 Mb (in total 4.4 Mb), is close to fixation in LWS (Figure 4). This haplotype was already close to fixation by generation 40, possibly indicating that selection has been relatively strong, and the resulting pattern resembles a hard sweep in LWS. Contrastingly, multiple haplotypes segregate in HWS, with a mixture of nucleotide positions that are divergently fixed when compared to LWS, intermixed with positions that segregate for both LWS and unique HWS alleles.

We observed a high level of divergence between HWS and LWS within this region, implying that the long haplotype fixed in LWS is not present in the HWS at generation 55 because it was selected out of this line. Highly dissimilar regions are intermixed with short, but continuous, stretches where the HWS and LWS haplotypes are nearly identical. Additionally, the boundaries between these shared and differentiated regions are distinct. They likely represent historical recombination events shared between one or more selected haplotypes, rather than being multiple classic hard sweeps, as these are expected to result in a gradual breakdown of population-wide linkage disequilibrium.

This interpretation is supported by other observations. First, for sharing of such interspersed haplotypes to be possible between the lines, formative recombination events must have occurred prior to the onset of bidirectional selection. Second, as several shared haplotype segments were sometimes observed in the divergent regions, such multiple events must have happened on the same haplotypes. Third, because the shared regions are often short (10s to a few 100 kb), the events are unlikely in a population with as small a population-size as the Virginia body weight chicken lines (effective population size, N_e_ ~ 35) (Marquez et al. 2010). Finally, if the recombination events occurred during the selection experiment, selection on the recombinant haplotypes must have been strong in order to only retain them in the lines. Such is unlikely given the highly polygenic genetic architecture of 8-week body weight and the dilution of selection pressure across the many loci. Therefore, these haplotype mosaics most likely represent recombinant founder haplotypes resulting from their population history, including the formation of the WPR, mild inbreeding, and finally intercrossing (Figure 1). Positive selection of this long haplotype to fixation in the LWS, coupled with negative selection for its removal from HWS, may serve as an explanation for why previous studies have seen a transgressive effect on 8-week body weight for SNP markers within this region. Zan et al. (2017) reported that while the HWS allele at the rs14916997 SNP marker (GGA1:169,408,309 bp) was associated with an increased 8-week body weight (additive genetic effect approximately 26 grams), the HWS allele at the nearby SNP marker rs316102705 (GGA1: 172,235,208 bp) was associated with a decrease in 8-week body weight (additive genetic effect of approximately −7 grams).

Notably, this region on chicken chromosome GGA1 appears often in association studies carried out in other populations, including comb traits (Shen et al. 2016), egg weight (Yi et al. 2015), feed intake (Yuan et al. 2015), abdominal fat percentage (Abasht and Lamont 2007; Sheng et al. 2013), shank metrics (Sheng et al. 2013), and growth and body weight at numerous life stages (Xie et al. 2012; Sheng et al. 2013; Abdalhag et al. 2015; Zhang et al. 2015). This may reflect a shared ancestral variant that has spread in domestic populations due to its beneficial effect, or that this is a gene rich region associated with many functionally important genes. Recently, an insertion (GGA1: 169,399,420) located upstream of the miR-15a-16 precursor was strongly associated with growth traits in an Xinghua & White Recessive Rock F_2_ cross, where presence of this variant results in an altered hairpin formation, reduction of miR-16 expression, and increased body weight, bone size, and muscle mass (Jia et al. 2016). Although this insertion was also present in multiple chicken breeds, and at high frequencies in broiler breeds (Jia et al. 2016), in our population, the insertion was present in LWS on the long, fixed haplotype, and was absence from the HWS, contrary to expectations that the insertion would substantially increase body weight.

This observation may be explained by alternative hypotheses; i.e. that this insertion: i) has an effect also in our lines but is linked to another polymorphism with a stronger opposite effect, ii) does not have an effect in our lines due to genetic background, iii) does not have an effect at all, suggesting that earlier reports revealed association between this polymorphism and the studied traits, rather than causation. Alternatively, as is often the case, the functional variant may lie outside coding regions. Nevertheless, further work within this QTL region will be required to fully characterized the functional variant responsible for the large difference in 8-week body weight in the Virginia body weight lines.

## Conclusions

Over the course of long-term bidirectional selection, the Virginia body weight lines have experienced significant changes impacting behavioral, neurological, metabolic, and developmental processes (Dunnington and Siegel 1996). With a concerted effort to understand the fundamental genomics underlying these changes, we have employed linkage and association mapping, selective-sweep analyses using high-density SNP data and pooled genome sequencing across generations of the selected lines, relaxed lines, and a derived advanced intercross line (Wahlberg et al. 2009; Johansson et al. 2010; Pettersson and Carlborg 2010; Besnier et al. 2011; Pettersson et al. 2011; Ahsan et al. 2013; Pettersson et al. 2013; Sheng et al. 2015; Brandt et al. 2016). It has become clear that the difference in body weight between the selected lines relies on small to moderate effects from many loci, with the most recent analysis revealing 20 contributing loci (Zan et al. 2017).

Pooled genome resequencing revealed distinct hallmarks of selection in regions that were highly differentiated in the lines after 55 generations of long-term experimental selection.

More often than not, the regions show the persistence of haplotypic diversity in one line, contrasted by fixation in the other. Despite selection acting on the same pool of standing genetic variants, the genomic signatures of selection thus resemble classic hard sweeps in one line, contrasted to a mosaic pattern of divergence in the other. This haplotype mosaic emerges from recombination of multiple divergent founder haplotypes, probably shaped by historical bottlenecks, crossbreeding, and inbreeding, which forms the standing genetic variation available in this selection experiment.

## Acknowledgements

For help with genome sequencing, we thank Uppsala University SNP&SEQ Technology Platform (Science for Life Laboratory), a national infrastructure supported by the Swedish Research Council (VR-RFI) and the Knut and Alice Wallenberg Foundation. For computational resources, we thank the Uppsala Multidisciplinary Center for Advanced Computational Science and the associated Next generation sequencing Cluster and Storage (UPPNEX) project, funded by the Knut and Alice Wallenberg Foundation and the Swedish National Infrastructure for Computing (SNIC; project id: b2015010). This work was supported by the Swedish Research Council for Environment, Agricultural Sciences and Spatial Planning (DNR 221-2013-450 to ÖC) and various sources at Virginia Tech. Lastly, individuals at Virginia Tech for a range of roles in producing and maintaining the chickens across more than half a century.

## Authors’ Contributions

ÖC and PBS designed the study. ML performed analyses. CFH and PBS designed the pedigrees and managed the Virginia body weight chickens, collected samples, and provided DNA. ML and ÖC wrote the manuscript. All authors read and approved the final manuscript.

